# A novel pathway of functional microRNA uptake and mitochondria delivery

**DOI:** 10.1101/2022.11.07.515397

**Authors:** Jiachen Liu, Weili Li, Jianfeng Li, Eli Song, Hongwei Liang, Weiwei Rong, Xinli Jiang, Nuo Xu, Wei Wang, Shuang Qu, Yujing Zhang, Chen-Yu Zhang, Ke Zen

## Abstract

Extracellular miRNAs serve as signal molecules in the recipient cells. Uptake of extracellular miRNAs by the recipient cells and their intracellular transport, however, remains elusive. Here we show RNA phase separation as a novel pathway of miRNA uptake. In the presence of serum, synthetic miRNAs rapidly self-assembly into ∼110nm discrete nanoparticles which enable miRNAs’ entry into different cells. Depleting serum cationic proteins prevents the formation of such nanoparticles and thus blocks miRNA uptake. Different from lipofectamine-mediated miRNA transfection in which the majority of miRNAs are in lysosomes of transfected cells, nanoparticles-mediated miRNA uptake predominantly delivers miRNAs into mitochondria in a polyribonucleotide nucleotidyltransferase 1-dependent manner. Functional assays further show that the internalized miR-21 via miRNA phase separation enhances mitochondrial translation of Cytochrome b, leading to increase in ATP and ROS reduction in HEK293T cells. Our findings reveal a previously unrecognized mechanism for uptaking and delivering functional extracellular miRNAs into mitochondria.

**Synopsis:** 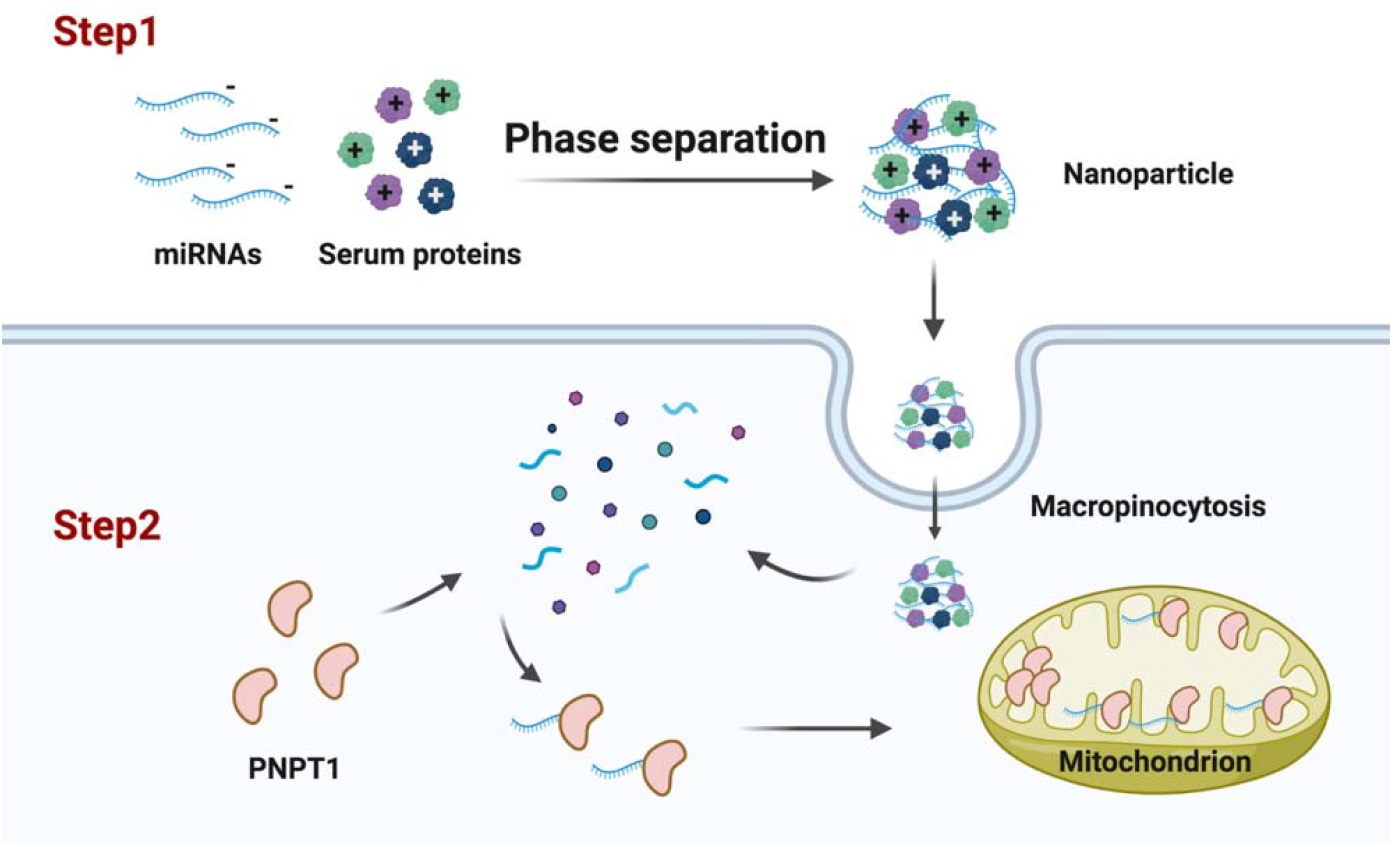

RNA phase separation-based extracellular miRNA uptake and PNPT1-mediated mitochondrial delivery of internalized miRNAs

- miRNAs can self-assembly into ∼110nm nanoparticles to enter various cells in the presence of serum
- miRNA phase separation is mediated by serum cationic proteins
- Internalized miRNAs via this nanoparticle pathway are predominantly delivered to mitochondria
- Mitochondrial delivery of the internalized miRNAs is mediated by PNPT1

## Introduction

Given their ability to interact with a broad, yet specific set of target genes through a base-pairing mechanism, microRNAs (miRNAs) significantly contribute to the gene expression (Zamore, 2002). It is widely accepted that almost all cell types can produce and secrete miRNAs. As the extracellular miRNAs released by host cells can enter the recipient cells where they serve as a novel class of signaling molecules to modulate the expression of target genes and the function of recipient cells (Kosaka *et al*, 2010; Zhang *et al*, 2010), miRNAs also play a critical role in horizontal gene regulation (Chim *et al*, 2008). The mechanisms of transferring functional miRNAs from host to recipient cells, however, remain elusive though the previous studies have illustrated that uptake of extracellular miRNAs by host cells depends on the supramolecular structure and composition of miRNA carriers (Syed *et al*, 2019), as well as the nature of recipient cells (Horibe *et al*, 2018).

As a ∼22nt hydrophilic molecule, miRNA is not able to directly cross cell surface membranes. Extensive studies demonstrate three major pathways for delivering extracellular miRNAs into the recipient cells: First, miRNA uptake by recipient cells is mediated by extracellular vesicles (EVs), particularly exosomes(Valadi *et al*, 2007). In response to various stimuli, cells can selectively pack miRNAs into exosomes and release them via exocytosis pathway (Zhang *et al*., 2010). Uptake of EV-encapsulated miRNAs by recipient cells can be carried out via endocytosis, phagocytosis, micropinocytosis or macropinocytosis depending upon the type of recipient cells (Chen *et al*, 2012; Tian *et al*, 2014; Xu *et al*, 2013). Besides the EV-encapsulated miRNAs, a significant portion of extracellular miRNAs are in various free-floating complexes associated with RNA-binding proteins, such as Argonaute 2 (AGO2) or nucleophosmin (NPM1)(Arroyo *et al*, 2011). Uptake of the extracellular miRNAs by recipient cells can be also mediated by these RNA-binding protein(Prud’homme *et al*, 2016; Wang *et al*, 2010). For instance, AGO2-bound miRNAs were reported to be taken up by recipient cells via neurophilin-1 (NRP1)(Prud’homme *et al*., 2016). Extracellular miRNAs can be also associated with highly abundant miRNA carriers such as high- and low-density lipoproteins (Michell & Vickers, 2016). Uptake of these lipoprotein-bound miRNAs via lipoprotein-mediated pathway was established by various investigators (Frank *et al*, 2019; Vickers *et al*, 2011). However, these three pathways described above have limitations and cannot explain the rapid uptake of extracellular miRNAs that are not encapsulated in EVs or associated with RNA-binding proteins or lipoproteins (Castanotto *et al*, 2016; Takahashi *et al*, 2017).

In the present study, we demonstrate a novel pathway of general rapid miRNA uptake in various cell types. Different from uptake mediated by EVs and other miRNA carriers, this miRNA uptake pathway is dependent upon RNA phase separation mediated by serum cationic proteins. More importantly, instead of accumulated at lysosomes for degradation, the internalized miRNAs through this unique pathway are mainly delivered into recipient cell mitochondria where they may modulate mitochondrial gene functions.

## Results

### Uptake of miRNA by various cells in the presence of serum and uniquely delivered into mitochondria

To explore the interaction between extracellular miRNAs and host cells, we incubated Hela cells with fluorescent 5’-Cy5-miR-29a (miR-29a attaching fluorophore Cy5 at the 5’-terminus) at 37°C in the presence of 10% fetal bovine serum (FBS). To our surprise, we observed that, instead of adherent to the cell surface, 5’-Cy5-miR-29a rapidly entered the cells in a time-dependent manner (Fig. 1a). Quantitative analysis of cell-associated fluorescence confirmed the time-dependent fluorescent miR-29a uptake by Hela cells (Fig. 1b). Uptake of miR-29a in Hela cells was confirmed using 5’-Cy3-miR-29a (miR-29a attaching fluorophore Cy3 at the 5’-terminus) (Extended Data, figure 1a), while incubating cells with fluorophore Cy5 alone displayed no cell-associated fluorescence (Extended Data, figure 1b). These results suggest that miR-29a uptake is independent of fluorophore modification. As expected, entry of miR-29a-Cy5 into HeLa cells was abolished at low temperature or after ATP depletion (Fig. 1, a and b), confirming that miRNA uptake by the recipient cells is an active process. However, uptake of 5’-Cy5-miR-29a by HeLa cells at 37°C was also completely blocked in the absence of serum (Fig. 1, a and b), suggesting that 5’-Cy5-miR-29a uptake by HeLa cells at 37°C is mediated by serum. The uptake of miRNAs in the presence of serum was further validated in different cell types including HEK293T, A549, U87MG, Min6 and SGC-7901, using fluorescent miRNAs (Extended Data, figure 1c). As disrupting cellular microfilament networks can block endocytosis and micropinocytosis (Al Soraj *et al*, 2012; da Costa Goncalves *et al*, 2021), we further analyze the uptake of 5’-Cy5-miR-29a in the presence of serum after treating Hela cells with cytochalasin B, rottlerin, dynarose, nocodazole or other cytoskeleton disrupting reagents. After 1 h incubation, we found that rottlerin and nocodazole, two macropinocytosis inhibitor (Hufnagel *et al*, 2009), strongly prevented cellular uptake of 5’-Cy5-miR-29a, whereas other reagents displayed no inhibition on 5’-Cy5-miR-29a uptake (Fig. 1c, Extended Data, figure 1d). This result suggests that uptake of 5’-Cy5-miR-29a is likely via the pathway of macropinocytosis, which is in agreement with the notion that miRNA uptake in the presence of serum is an active process and dependent upon dynamic arrangement of the cellular microfilament network.

**Figure 1.**
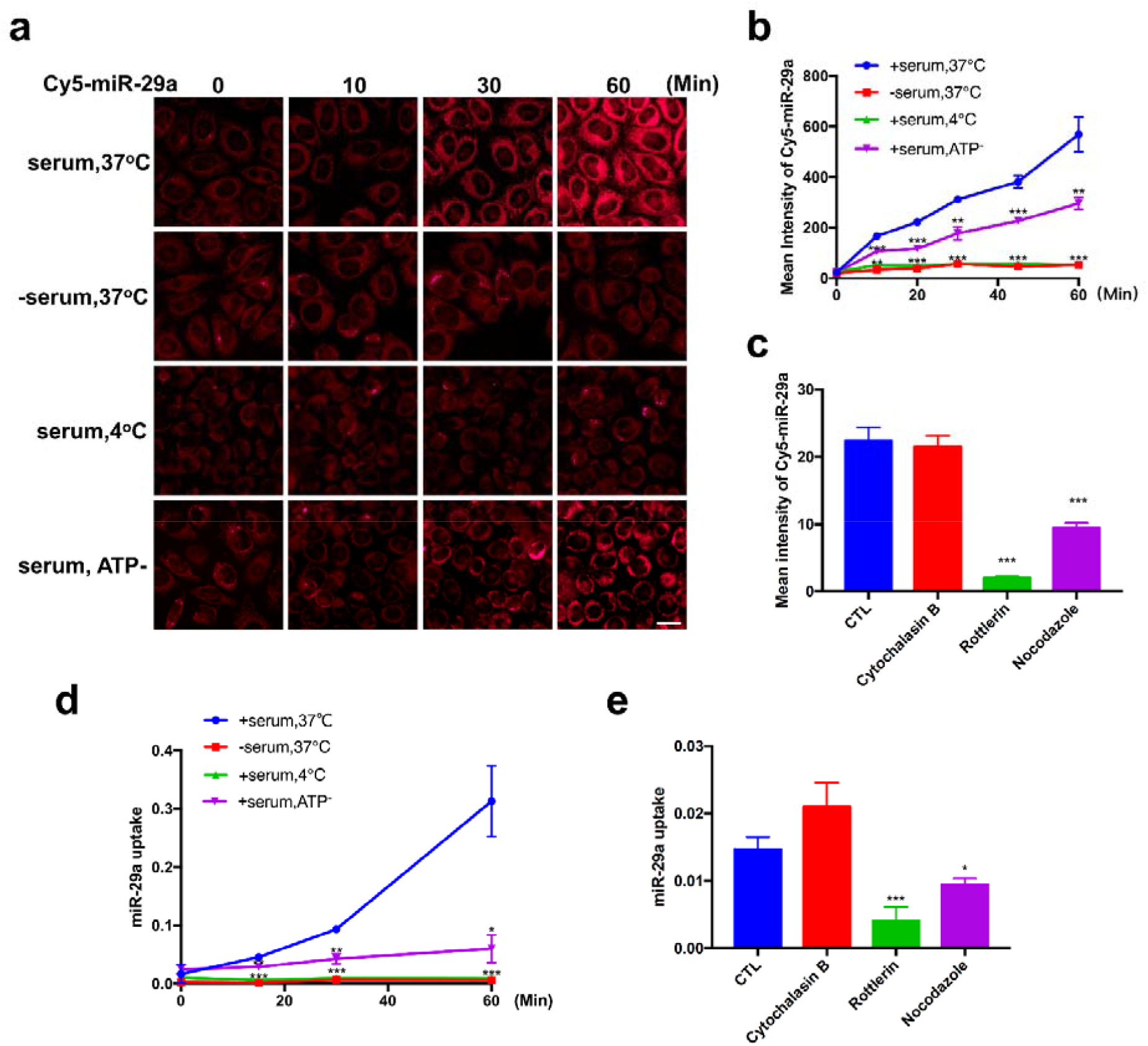
Direct uptake of extracellular miRNAs by recipient cells in the presence of serum. **a**, Time course of Cy5-miR-29a uptake by HeLa cell under various conditions. Scale bar, 20μm. **b**, Quantification of Cy5-miR-29a uptake by HeLa cells with various treatment. **c**, Inhibition of Cy5-miR-29a uptake by rottlerin and nocodazole. **d**, Time-dependent miR-29a uptake by HeLa cells quantified by qRT-PCR under various conditions. **e**, Inhibition of miR-29a uptake by Rottlerin and Nocodazole. The above experiments were repeated 3 times and samples at each time point of qRT-PCR assay were triplicated. *P<0.05, **P<0.01, ***P<0.001.

The uptake of extracellular miRNA in the presence of serum was further validated using synthetic miR-29a. As shown by qRT-PCR analysis (Fig. 1d-e), miR-29a was rapidly taken up by HeLa cells at 37°C in the presence of serum, whereas the time-dependent miR-29a uptake was abolished in the absence of serum, low temperature or ATP depletion (Fig. 1d). Similarly, blockade of macropinocytosis process by rottlerin markedly blocked miR-29a uptake (Fig. 1e).

Next we examined the intracellular location of the internalized 5’-Cy5-miR-29a. In this experiment, HeLa cells were incubated with 5’-Cy5-miR-29a in the presence of 10% FBS 37°C for 15 min, and then washed off the unbound 5’-Cy5-miR-29a and continuously incubated for 60 min. As shown in Fig. 2a, 5’-Cy5-miR-29a mainly located around the peripheral area inside cells after 15 min incubation, demonstrating a rapid internalization of 5’-Cy5-miR-29a. Strikingly, following 60 min and 120 min tracing, the internalized 5’-Cy5-miR-29a was predominantly transported to mitochondria-like subcellular structures. Double staining subcellular structures with specific markers confirmed that internalized 5’-Cy5-miR-29a was primarily delivered to mitochondria (mito) not lysosomes (lyso) or endoplasmic reticulum (ER) (Fig. 2b). To validate the fluorescent miR-29a tracing result, we incubated Hela cells with hcmv-miR-UL148D, an exogenous miRNA, in the presence of 10% FBS for 0, 1 and 6 h, and then examined the subcellular distribution of hcmv-miR-UL148D by qRT-PCR. As shown in Fig. 2c, levels of hcmv-miR-UL148D in whole cell and mitochondria were time-dependently increased, and the majority of internalized hcmv-miR-UL148D was detected in mitochondria after 6 h incubation.

**Figure 2.**
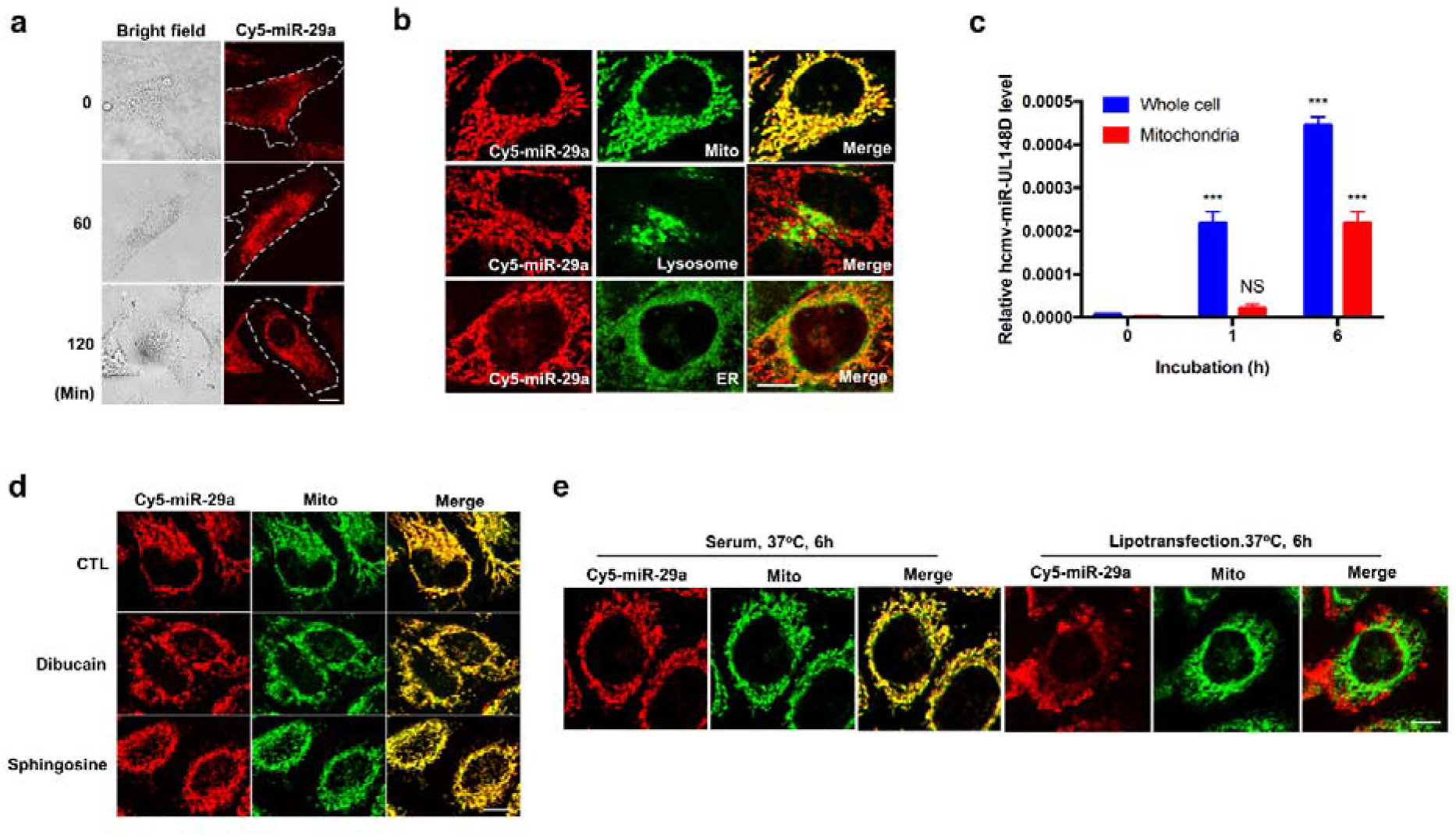
Selective delivery of internalized fluorescent miRNAs into mitochondria of recipient cells. **a**, Tracing the internalized Cy5-miR-29a in HeLa cells for 2 h after 15 min uptake. **b**, Distribution of the internalized Cy5-miR-29a at mitochondria but not lysosome and ER. **c**, qRT-PCR analysis of exogenous hcmv-miR-UL148D in whole cell lysate and mitochondria at different time points of incubation. **d**, Distribution of the internalized Cy5-miR-29a at mitochondria after disrupting mitochondrial membrane potential by Dibucaine and Sphingosine. **e**, Comparison of intracellular localization of Cy5-miR-29a via incubating cells with Cy5-miR-29a in the presence of 10% FBS and direct Lipotransfection of fluorescent miRNA in HeLa cells at 6 h post-treatment. Scale bars, 10μm. The above experiments were repeated 3 times and samples in in each time point of qRT-PCR assay were triplicated. NS, no significance. ***P<0.001

Given that alteration of mitochondria membrane potential can change mitochondria shape and localization(Umbaugh *et al*, 2021), we examined the intracellular localization of 5’-Cy5-miR-29a following disruption of mitochondria membrane potential by Dibucaine and Sphingosine(Heit *et al*, 2011). As shown in Fig. 2d, treatment with Dibucaine and Sphingosine did not alter the co-localization of internalized 5’-Cy5-miR-29a with mitochondria tracker in HeLa cells. The localization of internalized 5’-Cy5-miR-29a in the presence of 10% FBS was also compared to that through lipofectamine transfection (Fig. 2e). Different from lipofectamine transfection in the absence of serum, in which most 5’-Cy5-miR-29a was observed in lysosomes at 6 h post-transfection, miRNA uptake in the presence of serum mainly delivered the internalized 5’-Cy5-miR-29a into mitochondria after 6 h incubation at 37°C. To exclude the possibility that mitochondria delivery of internalized miRNA via serum-mediated uptake may be restricted to certain miRNA or cell types, we tested various kinds of cell lines using different small RNA species such as miR-21-5p, miR-1-3p mimics, miR-122, hcmv-miR-UL148D and double-strand miR-29a (Sequences listed in Extended data, table 1). We found that, without exception, all these internalized small RNAs were delivered into mitochondria in different cells (Extended Data, figure 2a-b).

### Uptake of miRNA is dependent upon RNA phase separation mediated by serum cationic proteins

As uptake of miRNA requires the presence of serum, our initial hypothesis was that certain serum proteins might serve as miRNA receptor(s) or carriers for mediating the miRNA uptake by the recipient cells. However, when we denatured the serum proteins by heat treatment (60°C, 1 h) prior to the incubation of Hela cells with exogenous hcmv-miR-UL148D, we found that hcmv-miR-UL148D uptake by HeLa cells was not affected by serum denature (Extended Data, figure 3). As previous studies suggest that RNA phase separation, such as forming nanoparticles, may facilitate the entry of RNA into cells(Beddoes *et al*, 2015), we examined whether miRNA phase separation occurred in the presence of serum. In this experiment, 50pmol/mL synthetic miR-29a was mixed with 10% FBS in DMEM and miRNA phase separation was monitored using Nanosight. Prior to this experiment, FBS was ultracentrifuged at 120,000g for 2 h and filtered using a 100kD cutoff ultra-filtration tube (Fig. 3d, left) at 4000g for 20 min to remove various microvesicles (MVs)(Skog *et al*, 2008; Valadi *et al*., 2007). The Nanosight results clearly indicated that miR-29a rapidly formed numerous ∼110 nm nanoparticles in the presence of 10% MV-free FBS, whereas no nanoparticles were detected in the DMEM containing miR-29a or MV-free serum alone (Fig. 3a-b). The formation and size of miR-29a nanoparticles in the presence of 10% MV-free serum were further analyzed by transmission electron microscope (TEM), which confirmed the formation of ∼110 nm nanoparticles after mixing miR-29a with 10% MV-free FBS (Fig. 3c). To test whether miRNA uptake relies on the formation of miRNA nanoparticles in the presence of 10% MV-free FBS, we separated the miR-29a in nanoparticles (NP-miR-29a) from ‘free’ miR-29a using centrifugation with a 100kD cutoff filter. The mixture of miR-29a with MV-free 10% FBS in DMEM was uploaded into the column, which was subjected to centrifugation (4000g, 20 min). The ‘free’ miR-29a and majority of serum proteins were filtered through and collected at the bottom fraction while the NP-miR-29a was harvested in the top chamber. As shown in Fig. 3d, right, the qRT-PCR assay showed that miRNA nanoparticles were rapidly formed after mixing miR-29a with 10% MV-free FBS, and more specifically, over 80% miR-29a was associated with nanoparticles.

**Figure 3.**
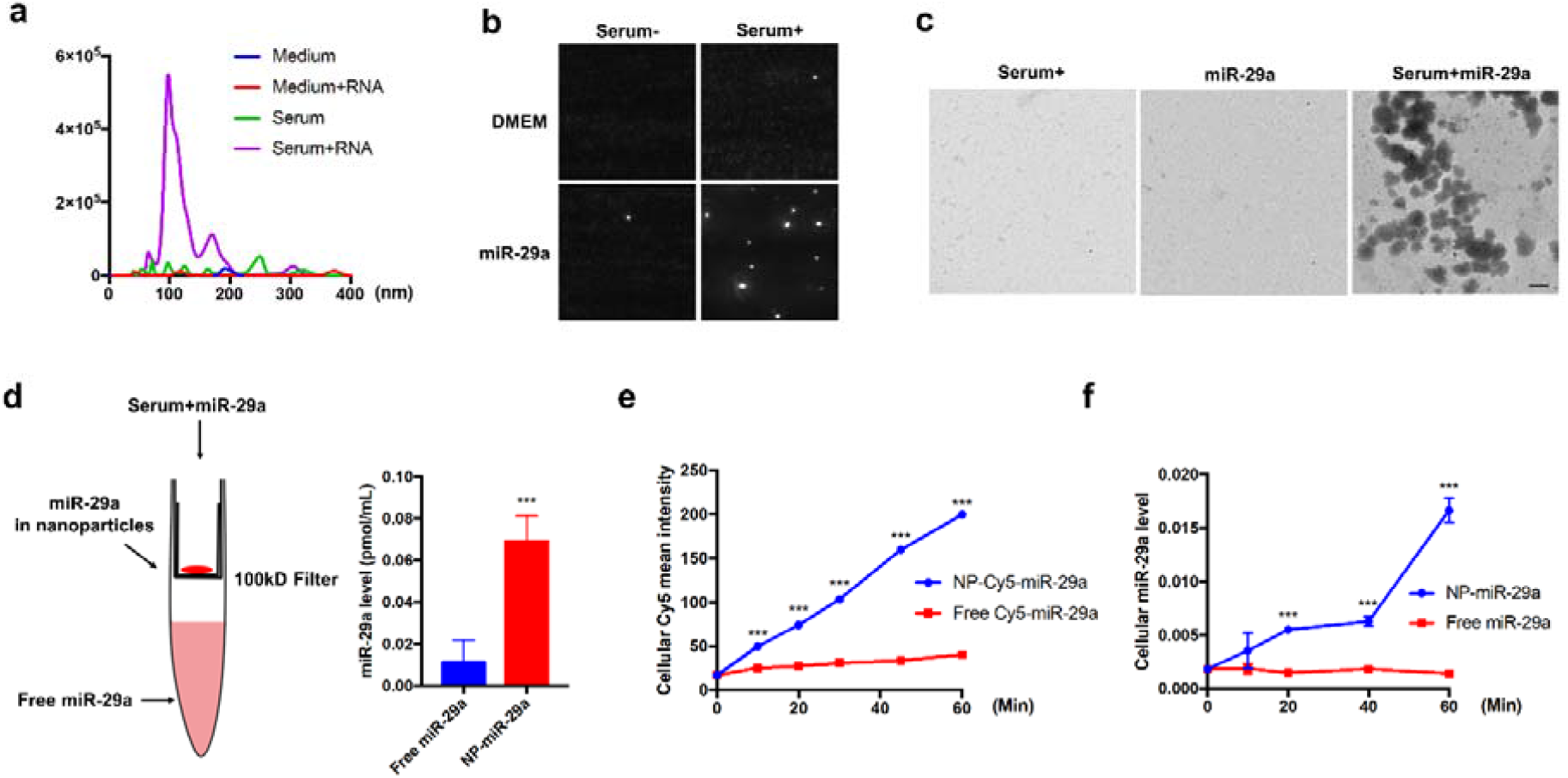
Uptake of extracellular miRNA in the presence of serum is dependent upon the formation of miRNA nanoparticles. **a-b**, Formation of ∼110nm miR-29a nanoparticles after mixing with 10% MV-free FBS assessed by Nanosight. **c**, TEM images of 10% MV-free serum, miR-29a and mixture of miR-29a and 10% MV-free serum. **d**, Schematic description of separating free miR-29a (free miR-29a) from miR-29a in nanoparticles (NP-miR-29a) via centrifugation with a 100kD cut-off filter. **e**, Uptake of free Cy5-miR-29a or NP-Cy5-miR-29a by HeLa cells assessed by fluorescence imaging. **f**, Uptake of free miR-29a or NP-miR-29a by HeLa cells assessed by qRT-PCR. The above experiments were repeated 3 times and samples at each time point in qRT-PCR assay were triplicated. Scale bars, 200nm. ***P<0.001.

We next investigated whether the nanoparticle formation in the presence of serum serves as the mechanism by which miRNAs were taken up by the recipient cells. In this experiment, both 5’-Cy5-miR-29a and hcmv-miR-UL148D were used. Various miRNAs were mixed with 10% MV-free FBS in DMEM and then separated into the nanoparticle fraction and the nanoparticle-free fraction, respectively. HeLa cells were then incubated with 5’-Cy5-miR-29a or hcmv-miR-UL148D in nanoparticle and nanoparticle-free fractions at 37°C for various time points, and the miRNA uptake by HeLa cells was assessed via fluorescence imaging system and qRT-PCR assay, respectively. As shown in Fig. 3e, left, cell-associated fluorescence quantification indicated that only 5’-Cy5-miR-29a in nanoparticle fraction entered the cells and uptake of 5’-Cy5-miR-29a by HeLa cells was in a time-dependent manner. No uptake of 5’-Cy5-miR-29a in nanoparticle-free fraction was observed in HeLa cells. Similar results were obtained in the uptake of hcmv-miR-UL148D assayed by qRT-PCR (Fig. 3e, right). As shown, the hcmv-miR-UL148D in nanoparticle fraction but not nanoparticle-free fraction was time-dependently internalized into HeLa cells.

Previous studies have shown that cationic proteins or peptides such as polyarginine (R9) are involved in formation of RNA nanoparticles (Dana Maria Copolovici, 2014; Wang *et al*, 2021). Given that serum contains various cationic proteins derived from basophil and eosinophils (Plager *et al*, 2006), we next tested whether the cationic proteins in serum are responsible for miRNA phase separation. As shown in Fig. 4a, positively charged proteins in MV-free FBS were depleted using ion exchange chromatography. As expected, MV-free FBS with cationic protein depletion failed to mediate miRNA phase separation. When miR-29a was mixed with 10% MV-free FBS in which cationic proteins had been depleted, no nanoparticles of miR-29a were detected (Fig. 4b). In line with our previous observation that denatured serum was able to mediate miRNA uptake (Extended Data, figure 3), mixing hcmv-miR-UL148D with 10% heat-treated MV-free FBS (60°C, 1 h) also formed hcmv-miR-UL148D nanoparticles at a size of ∼110 nm (Fig. 4c). Supporting the notion that the formation of RNA nanoparticles in the presence of serum is the key for miRNA uptake by recipient cells, depleting cationic proteins in FBS prevented the formation of miRNA nanoparticles, thus completely abolished the serum-mediated uptake of 5’-Cy5-miR-29a and hcmv-miR-UL148D by HeLa cells assessed by fluorescence imaging and qRT-PCR (Fig. 4d), respectively.

**Figure 4.**
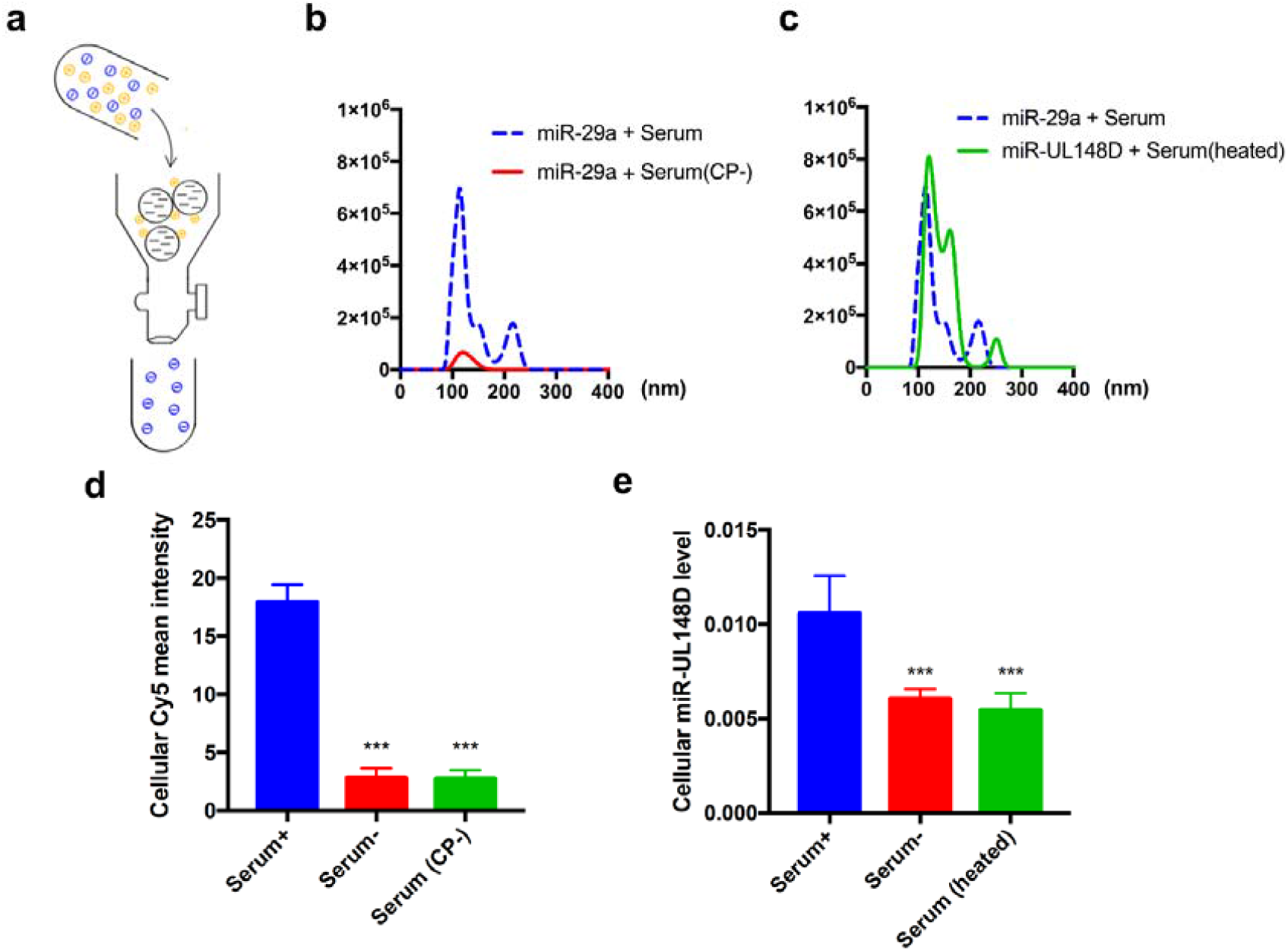
Formation of small RNA nanoparticles is mediated by serum cationic proteins. **a**, Schematic description of depletion of serum cationic proteins via ion-exchange column. **b**, Depletion of serum cationic proteins completely blocked the formation of miR-29a nanoparticles (red line). **c**, Denature of serum proteins via heating treatment (60°C) did not affect the formation of small RNA nanoparticles (green line). **d**, Uptake of Cy5-miR-29a or hcmv-miR-148D by HeLa cells in the presence of 10% MV-free FBS or 10% cationic protein-depleting MV-free FBS, serum (CP-), assessed by direct fluorescent imaging (left) and qRT-PCR (right), respectively. The above experiments were repeated 3 times and samples in each time point of qRT-PCR assay were triplicated. ***P<0.001.

### PNPT1 mediates the delivery of internalized miRNA to mitochondria of recipient cells

Recent studies have established PNPT1 as a transporter of various RNAs from the cytoplasm to mitochondria (Dhir *et al*, 2018; Wang *et al*, 2012). To test whether PNPT1 is involved in delivering the internalized miRNA via the miRNA phase separation pathway into mitochondria, we first generated HeLa cells that permanently expressed EGFP-PNPT1. To trace the internalized 5’-Cy5-miR-21, EGFP-PNPT1-expressing cells were incubated with 5’-Cy5-miR-21 at 37°C for 15 min in the presence of serum, and then washed with DMEM to remove unbound 5’-Cy5-miR-21. Cells were then continuously incubated for various time points. As shown in Fig. 5a, along with the incubation, more and more internalized 5’-Cy5-miR-21 (red) was co-localized with PNPT1 (green), implicating that PNPT1 may bind to the internalized 5’-Cy5-miR-21 and deliver it to mitochondria. In line with this, immunoprecipitating the internalized miR-21 tagged with biotin (biotin-miR-21) indicated its association with PNPT1. In the experiment, cells were incubated with biotin-miR-21 for 12 h in the presence of serum, and then subjected to cell lysis and immunoprecipitation of biotin-miR-21. Western blot analysis showed that significantly more PNPT1 was pulled down by biotin-miR-21 compared to the control (Fig. 5b).

**Figure 5.**
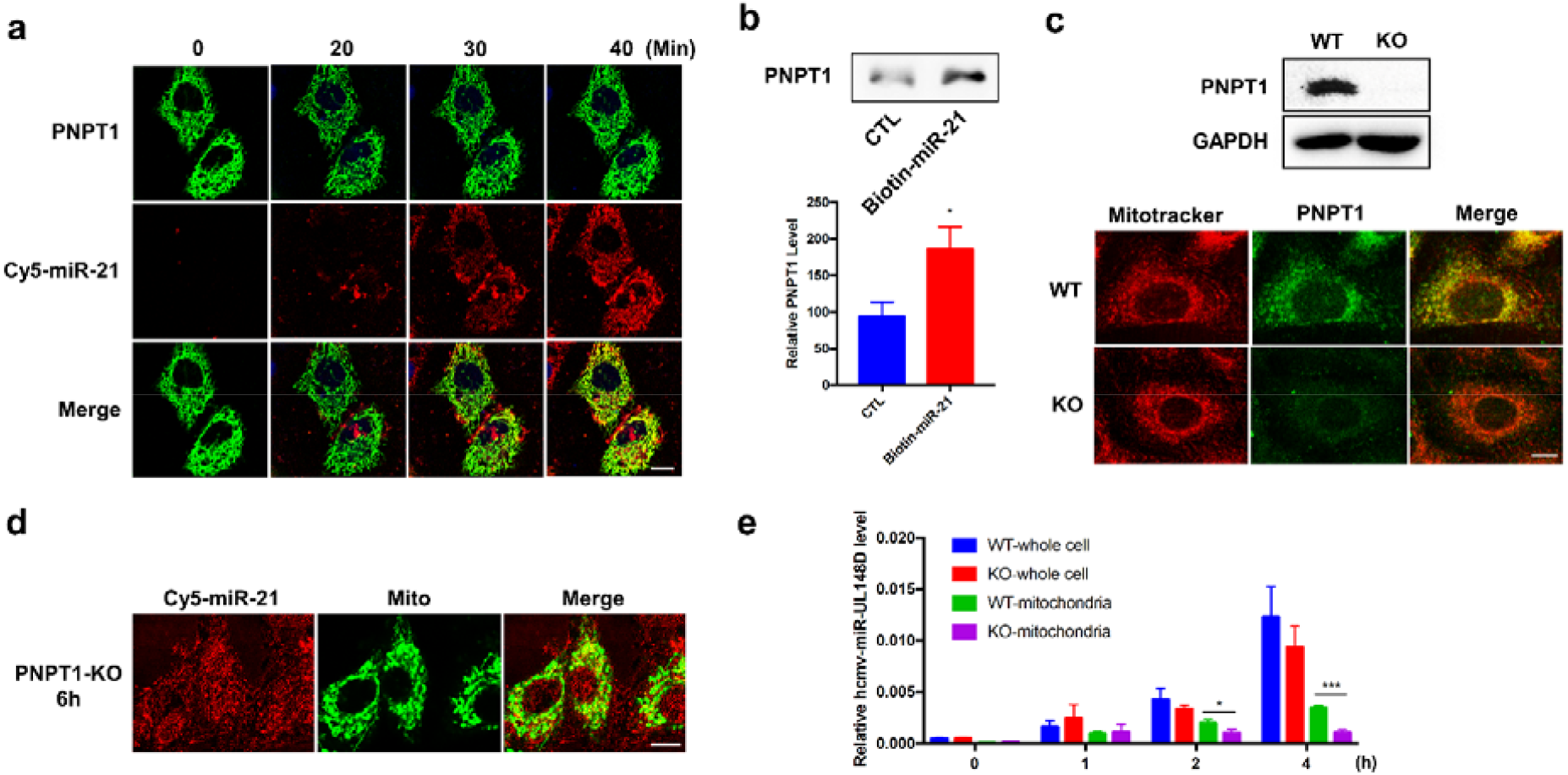
PNPT1 delivers the internalized miRNAs to mitochondria of recipient cells. **a**, Co-localization of internalized Cy5-miR-21 (red) with EGFP-PNPT1 (green) in HeLa following the uptake. **b**, Association of internalized biotin-miR-21 with PNPT1 in HEK293T cells assessed by immunoprecipitation of cellular biotin-miR-21. **c**, Generation of a permanent PNPT1-KO A549 cell line, and PNPT1 knockout in A549 cells was confirmed by Western blot analysis (upper) and immunofluorescence labeling (lower). **d**, Cy5-miR-21 failed to be delivered to mitochondria in the PNPT1-KO A549 cells after 6 h incubation, whereas its early uptake by PNPT1-KO A549 cells remained intact. **e**, Cellular distribution of exogenous hcmv-miR-UL148D in WT and PNPT1-KO A549 cells after incubation for various time points. Scale bars, 10μm. The above experiments were repeated 3 times and samples in each time point of qRT-PCR assay were triplicated. *P<0.05,***P<0.001.

To define the role of PNPT1 in mitochondria delivery of internalized miRNA, we generated a PNPT1-deficient A549 cell line using the CRISPR-Cas9 system. As shown in Fig. 5c, Western blot (top) and immunofluorescence analysis (bottom) confirmed that PNPT1 was successfully knocked out in A549 cells. Uptake and intracellular localization of 5’-Cy5-miR-21 clearly showed that the mitochondria delivery of internalized 5’-Cy5-miR-21 was blocked in PNPT1-KO A549 cells compared to that in WT A549 cells (Fig. 5d), though the earlier internalization process of 5’-Cy5-miR-21 in PNPT1-KO A549 cells was not affected. Mitochondria isolation and qRT-PCR assays further validated the role of PNPT1 in delivering the internalized miRNA to mitochondria. In this experiment, control A549 and PNPT1-KO A549 cells were incubated with hcmv-miR-UL148D in DMEM containing 10% FBS. After incubation for 0, 1, 2, and 4 h, cells were washed off, harvested and subjected to mitochondria isolation. After extraction of small RNAs, the level of hcmv-miR-UL148D in both mitochondria and mitochondria-depleted cellular fractions was assessed by qRT-PCR. As shown in Fig. 5e, hcmv-miR-UL148D was almost equally internalized into control A549 and PNPT1-KO A549 cells after short incubation (<1 h); however, the distribution of hcmv-miR-UL148D in two cell types was significantly different after 2 h incubation. In control A549 cells, the majority of internalized hcmv-miR-UL148D was detected in mitochondria after 4 h incubation. In contrast, only a small fraction of internalized hcmv-miR-UL148D was detected in mitochondria from PNPT1-knockdown A549 cells after 4 h incubation. Taken together, these results suggest that PNPT1 deficiency disrupts the transport of internalized miRNAs into mitochondria though it has no impact on the uptake of miRNA nanoparticles by the recipient cells.

### The internalized miR-21 upregulates mitochondrial cytochrome b translation in HEK293T cells

As our results show that intact miRNAs can enter the cells via phase separation in the presence of serum and subsequently delivered into mitochondria by PNPT1, we next ask whether these miRNAs are functionally active. In this experiment, HEK293T cells were incubated with miR-21 mimics in the presence of 10% FBS for various time points, and then the level and function of cytochrome b (CYB), a target gene of miR-21 (Li *et al*, 2016), was examined. As expected, fluorescence imaging of the internalized 5’-Cy5-miR-21 (Fig. 6a) showed that miR-21 was predominantly delivered into mitochondria of HEK293T cells. Served as a control, siRNA against CYB (si-CYB, sequence listed in Extended data, table 1.) was used to knock down CYB expression. HEK293T cells were incubated with si-CYB in the same way to miR-21 mimic for uptake. Western blot analysis of isolated mitochondria indicated that mitochondria delivery of miR-21 mimics increased mitochondrial CYB level (Fig. 4b), which is in agreement with a previous finding that miR-21 enhances translation efficiency of CYB in mitochondria(Li *et al*., 2016). As a negative control, incubating HEK293T cells with CYB siRNA in the presence of 10% FBS resulted in a marked CYB reduction.

**Figure 6.**
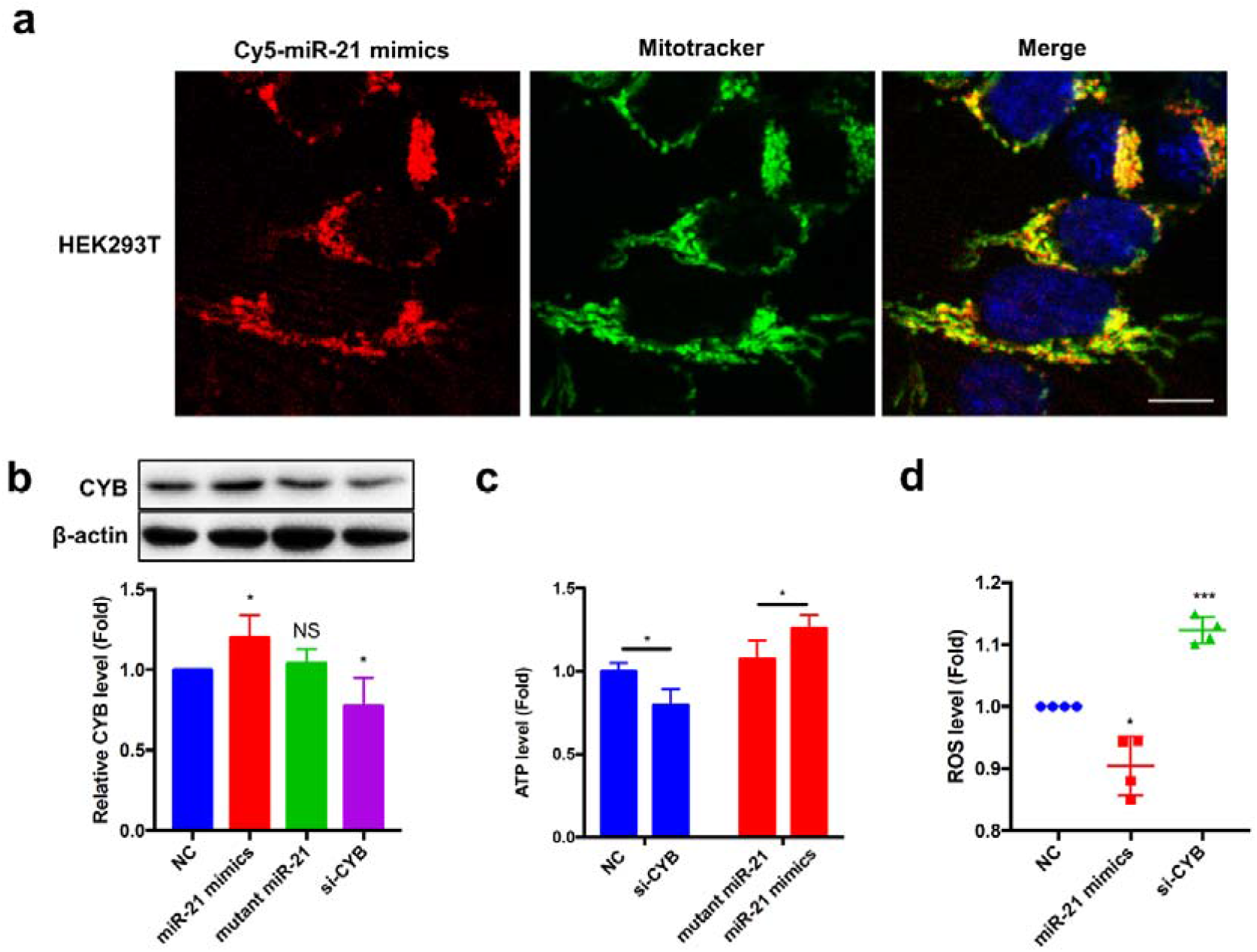
The internalized miR-21 increases mitochondrial cytochrome b level in the recipient cells. **a**, Delivery of the internalized miR-21 mimics to mitochondria in HEK293T cells after 6 h uptake mediated by the miRNA nanoparticle pathway. **b**, Upregulation of mitochondrial CYB by the internalized miR-21 mimics not the miR-21 mutant. **c**, ATP production in HEK293T cells following the uptake of miR-21 mimic, miR-21 mutant, si-CYB or control oligonucleotide (NC). **d**, Mitochondrial ROS level in HEK293T cells following uptake of miR-21 mimic, si-CYB or NC. In b-d, the experiments were repeated 4 times and samples in each time point were triplicated. Scale bars, 10μm. NS, no significance. *P<0.05. ***P<0.001.

As an essential component of respiratory complex III in mitochondria, CYB is positively associated with the production of cellular ATP (Cartron *et al*, 2014). Given that mitochondria CYB was upregulated or reduced by incubating cells with miR-21 mimic or CYB siRNA in the presence of serum, respectively, we next examined whether the cellular ATP level was altered accordingly. As shown in Fig. 6c, the cellular ATP level in HEK293T cells was increased by miR-21 but decreased by CYB siRNA. A previous study also reported that mitochondria CYB was involved in the reduction of cellular ROS (Cartron *et al*., 2014). We next measured the cellular ROS level following incubation with miR-21 mimics or CYB siRNA. The results showed that CYB siRNA markedly increased cellular ROS levels whereas miR-21 mimics decreased cellular ROS levels (Fig. 6d). These results collectively suggest that internalized miR-21 via serum-mediated miRNA phase separation pathway is intact and functionally active in the mitochondria of recipient cells.

## Discussion

In the present study, we demonstrate a novel RNA phase separation-mediated pathway for miRNA uptake, and that following this unique uptake process, the internalized miRNAs are delivered predominantly into mitochondria via PNPT1. In specific, extracellular miRNA forms ∼110nm discrete nanoparticles through interaction with serum cationic proteins. These miRNA nanoparticles can rapidly enter recipient cells through macropinocytosis. After internalization, the miRNAs bind to PNPT1, which delivers miRNAs into mitochondria.

Our results described a previously unrecognized uptake pathway for extracellular small RNAs by various cells, in which the formation of RNA phase separation in the presence of serum cationic protein is the key. The formation of RNA nanoparticles in the presence of protein or peptide with positive charge had been reported previously. Although the mechanism remains unknown, previous study showed that RNA drug could form large particles (∼500 nm) after mixing with a positively charged polyarginine peptide (R9), which facilitated the entry of RNA drugs into the recipient cells (Kumar *et al*, 2008). Different from the size of small RNA nanoparticles we observed in this study, the large size of RNA particles they detected may be due to the higher concentration and strong positive charge of the peptide. Indeed, when the ratio of R9 peptide vs RNA decreased, the size of RNA microparticles reduced (Law *et al*, 2008). Our data also suggest that the positive charge of serum cationic proteins but not protein themselves controls the miRNA phase separation since depleting cationic proteins in serum prevented the formation of miRNA nanoparticles whereas serum protein denature failed to affect miRNA nanoparticle production.

Different from simply membrane fusion, this unique uptake mediated by small RNA phase separation is an active process, which requires ATP and strong membrane fluidity. Depleting intracellular ATP or decreasing temperature strongly blocked miRNA uptake. In line with this, such RNA nanoparticle-mediated miRNA uptake required a dynamic arrangement of cellular microfilament network. We found that rottlerin strongly prevented the internalization of 5’-Cy5-miR-29a (Fig. 1c; Extended Data, figure 1d), suggesting that miRNA uptake via small RNA nanoparticles is through a mechanism similar to micropinocytosis (Sarkar *et al*, 2005). Previous studies have demonstrated that extracellular miRNA uptake by mammalian cells through several mechanisms, including exosome-mediated endocytosis or micropinocytosis/ micropinocytosis, and the internalization process mediated by various RNA-binding proteins (Frank *et al*., 2019; Prud’homme *et al*., 2016; Vickers *et al*., 2011; Wang *et al*., 2010). Given that small RNA phase separation mediated by serum cationic protein is not dependent upon the specific RNA receptors or binding proteins, miRNA uptake mediated by RNA nanoparticles may serve as a general pathway for various small RNAs, especially those not encapsulated in exosomes or associated with RNA-binding protein or lipoproteins (Castanotto *et al*., 2016; Takahashi *et al*., 2017).

Different from the cell Lipotransfection for miRNA delivery, in which the majority of intracellular miRNAs were accumulated at lysosomes, the internalized miRNAs via small RNA nanoparticles were mainly sorted into mitochondria. This finding suggests that mitochondria may serve as an important subcellular organelle for the storage of extracellular miRNAs, especially the miRNAs internalized into the recipient cells via the macropinocytosis pathway. Through co-localization analysis and gene silence assay, we identified mitochondria-associated PNPT1 as a transporter for delivering internalized miRNA into mitochondria. As a RNA-binding protein, PNPT1 plays multiple role in RNA transport and degradation (Wang *et al*., 2012). PNPT1 has been shown to be involved in transport of various RNAs including miRNAs to mitochondria, in which it controls mitochondria RNA homeostasis (Sokhi *et al*, 2013) and also prevents the leakage of mitochondria double-stranded RNAs into the cytoplasm (Dhir *et al*., 2018). In line with this, our results show that PNPT1 is responsible for selectively delivering the internalized miRNAs via RNA nanoparticle-mediated uptake pathway to mitochondria. Although PNPT1 also possesses activity in cleaving single stranded RNA (Cameron *et al*, 2019), both qRT-PCR and functional assay in the present study suggest that at least some extracellular miRNAs internalized through small RNA phase separation mechanism are intact, and can regulate the expression of target genes in mitochondria. In summary, our studies reveal a general extracellular miRNA uptake pathway based on RNA phase separation, and a selective transport of these internalized miRNA into mitochondria mediated by PNPT1.

## Materials and Methods

### Reagents, cell lines and culture conditions

HeLa, A549, HEK293T, SGC-7901, MIN6 and U87MG cell lines were purchased from American Type Culture Collection (ATCC; Manassas, VA). SGC-7901 cells were cultured in RPMI-1640 (Gibco; Carlsbad, CA) supplemented with 10% (v/v) fetal bovine serum (FBS) (Genial; Brighton, CO). MIN6 cells were maintained in RPMI-1640 supplemented with 15% (v/v) FBS, 0.1% 2-hydroxy-1-ethanethiol (Sigma Aldrich; Saint Louis, MO). All other cells were cultured in DMEM (Gibco; Carlsbad, CA) supplemented with 10% (v/v) FBS at 37°C under 5% CO_2_.

### Antibodies and artificial Synthetic RNA molecules

Antibodies against GAPDH and β-actin, as well as goat anti-rabbit and goat anti-mouse IgG, were purchased from Cell Signaling Technology (Danvers, MA), Antibody against mt-CYB was obtained from Affinity Biosciences (Beijing, China). Antibody against PNPT1 was purchased from Santa Cruz Biotechnology (Santa Cruz, CA). The miRNAs, miRNA mimics, siRNAs, single-strand RNAs, double-strand RNAs and control RNAs were obtained via chemical synthesis (Biosyntech; Suzhou, China).

### A549 PNPT1-KO cell line construction

PNPT1 gene was knock out in A549 cell line using the CRISPR-Cas9 System. The gRNA: ATAGTGCTCGCACTTGCAAC (designed in http://crispr.mit.edu/) was linked into the CRISPR Cas9 Plasmid (Genscript; Nanjing, China) and transform into competent cell (Trelief 5a, Tsingke; Beijing, China). Harvest plasmids after cultivation were transfected into the cells using lipofectamine 3000 (Invitrogen; Carlsbad, CA) according to manufacture’s protocol. Cells were then screened out by 50μM Puromycin (Sangon Biotech; Shanghai, China) and seeded into 96 well plate to cultivate monoclonal cell. Subculture the monoclonal cells and then analyze the PNPT1 protein by Western blotting.

### Quantification of miRNAs in mitochondria and whole cell lysate

Cells cultured on 10 cm dishes were incubated with 50nM synthetic miRNAs in DMEM containing 10% MV-free FBS for a period time indicated before harvest. The cell culture medium was then replaced by 10% FBS DMEM with 10 mM RNase inhibitor RVC (Beyotime; Shanghai, China). Mitochondria were isolated using Mitochondria Isolation Kit (Beyotime; Shanghai, China). The purity of mitochondria and whole cell was determined by monitoring mitochondria marker TOM20 and cell marker β-actin (Extended Data, figure 4). Total RNAs in mitochondria or whole cell lysate were extracted by TRIzol (Sigma Aldrich; Saint Louis, MO). The level of miRNAs was analyzed by qRT-PCR on Roche Lightcycler 96 Detection System (Roche, Auckland, New Zealand) using a stem-loop RT probe (Applied Biosystems, Forster City, CA).

### Nanoparticle formation and separation

This experiment was performed on Nanosight NS300 (Malvern PANalytical, England) to test the diameter of particles. Prior to testing, serum was ultracentrifuged (120,000g, 4°C, 2 h) and filtered (using a 100 kD cutoff ultra-filtration filter, 4000g, 20 min) to deplete microvesicles (MV)(Skog *et al*., 2008; Valadi *et al*., 2007). The miRNA (50 nM) was then mixed with DMEM or DMEM containing 10% MV-free serum for 5 min at room temperature to form nanoparticles. To separate nanoparticle-associated miRNAs from free miRNAs following miRNA phase separation in the presence of 10% MV-free FBS, the miRNA-serum mixture was separated by centrifugation (4,000g, 20 min) with a 100kD cutoff filter. The miRNAs in nanoparticles (NP-miRNAs) were collected in the upper chamber while free miRNAs were collected in the lower part.

### Immunofluorescence assay

HeLa cells were transfected with PNPT1-EGFP-expressing plasmid 48 h prior to the incubation of cells with fluorescent miRNAs. HeLa cells were then incubated with 50 nM synthetic Cy5-miR-29 in DMEM supplemented with 10% FBS for 15 min, followed by washing off Cy5-miR-29a and replaced with new culture medium. Immunofluorescence imaging was obtained by Zeiss LSM 880 confocal microscope (Zeiss, Oberkochen, Germany). For immunofluorescence labeling of PNPT1 or marker proteins for mitochondria, lysosome or ER, cells were fixed by 4% paraformaldehyde (10 min, room temperature) followed by permeabilization and blocking using 5% BSA (Sangon Biotech; Shanghai, China).

### Ion exchange chromatography

Econo-Column (BioRad; Herculs, CA) and SP Sepharose Fast Flow (GE; Boston, MA) were used to deplete serum cationic proteins. After putting the beads into the column and balancing by 1M KCL, 10% MV-free FBS was added into the column. Gently mix for a while, serum with cationic protein depletion was collected in the outflow fraction.

### RNA immunoprecipitation

HEK293T cells were incubated with biotin-labeled miR-21 for 12 h before harvest. Then Streptomycin beads (MedChemExpress, NJ, USA) were added to the cell lysate to pull down the complex of biotin-miR-21 and associated proteins (4°C, 2 h). After 3 times washed by RIP buffer I (10 mM Tris-HCl, pH 7.5, 1 mM EDTA, 1 M NaCl, 0.01% Tween-20), proteins associated with biotin-miR-21 then eluted by RIPA lysate (Beyotime; Shanghai, China) 100°C 5min. PNPT1 level was finally tested by Western blot.

### Cell ATP detection and mitochondrial reactive oxygen species (mt-ROS) detection

Cellular ATP production was detected using ATP Assay Kit according to the manufacture’s protocol (Beyotime; Shanghai, China). To test the level of mitochondria ROS (mt-ROS), HEK293T cells were seeded into 24-well plate and incubated with synthetic negative control, miR-21 mimics, miR-21 mutant or CYB siRNA (si-CYB) for 48 h. The mt-ROS level was measured with MitoSox Red (Invitrogen, Carlsbad, CA) by flow cytometry.

## Data Avaliablity

All data and materials support the reported claims and comply with standards of data transparency. Data will be made available on reasonable request.

## Acknowledgements

The authors thank Ms. Jane Zen (Wellesley College, Boston, MA) for critical reading and constructive discussion of the manuscript.

## Author Contributions

K.Z, Y.Z, C.Z designed the experiments. J.L, W.L, J.L, A.S, H.L, W.R, X.J, N.X, W.W and S.Q performed the experiments and analyzed results. K.Z and J.L drafted the manuscript. All authors critically revised and approved the final version.

## Supporting Information

This work was supported by grants from the Ministry of Science and Technology of China (2018YFA0507100) and the National Natural Science Foundation of China (82170692, 31771666).

## Extended Data for

**Extended Data | Table 1.**
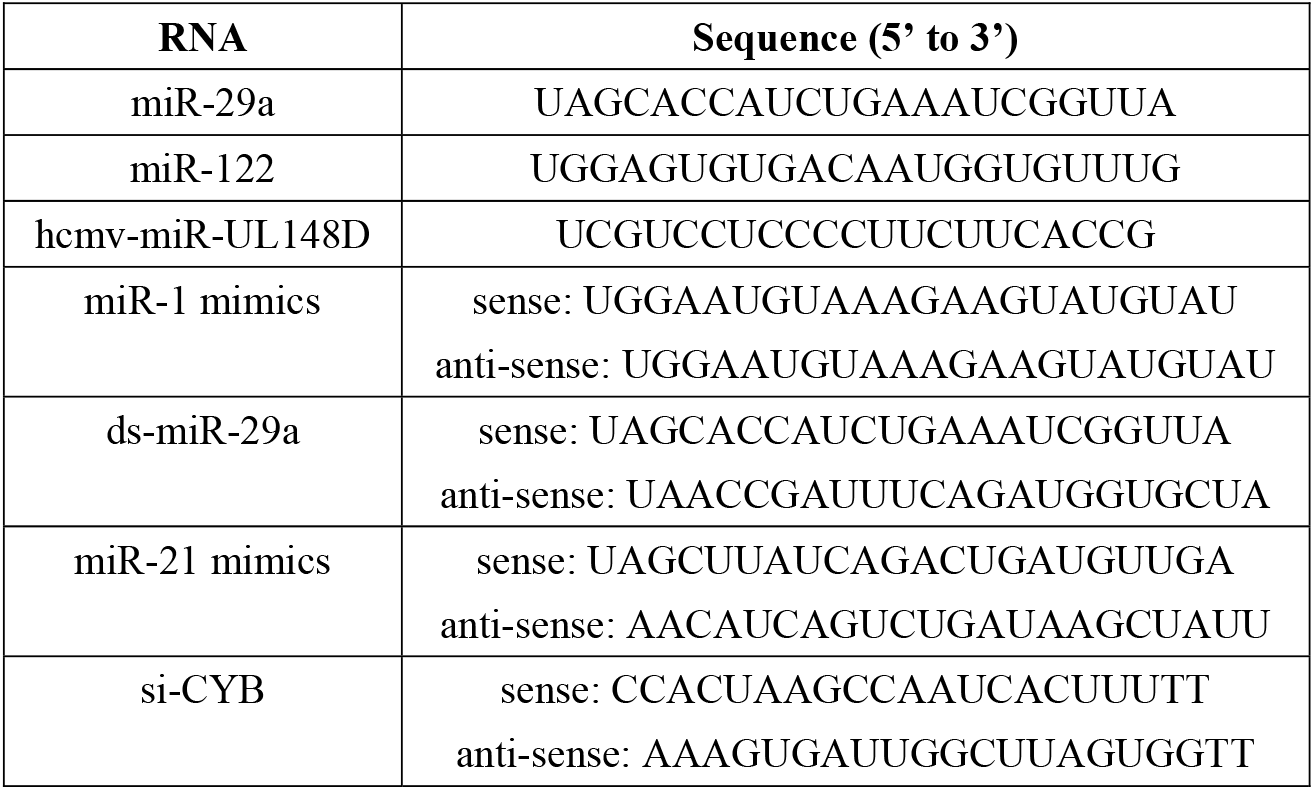
Sequence of RNAs used in this experiment.

**Extended Data | Figure 1.**
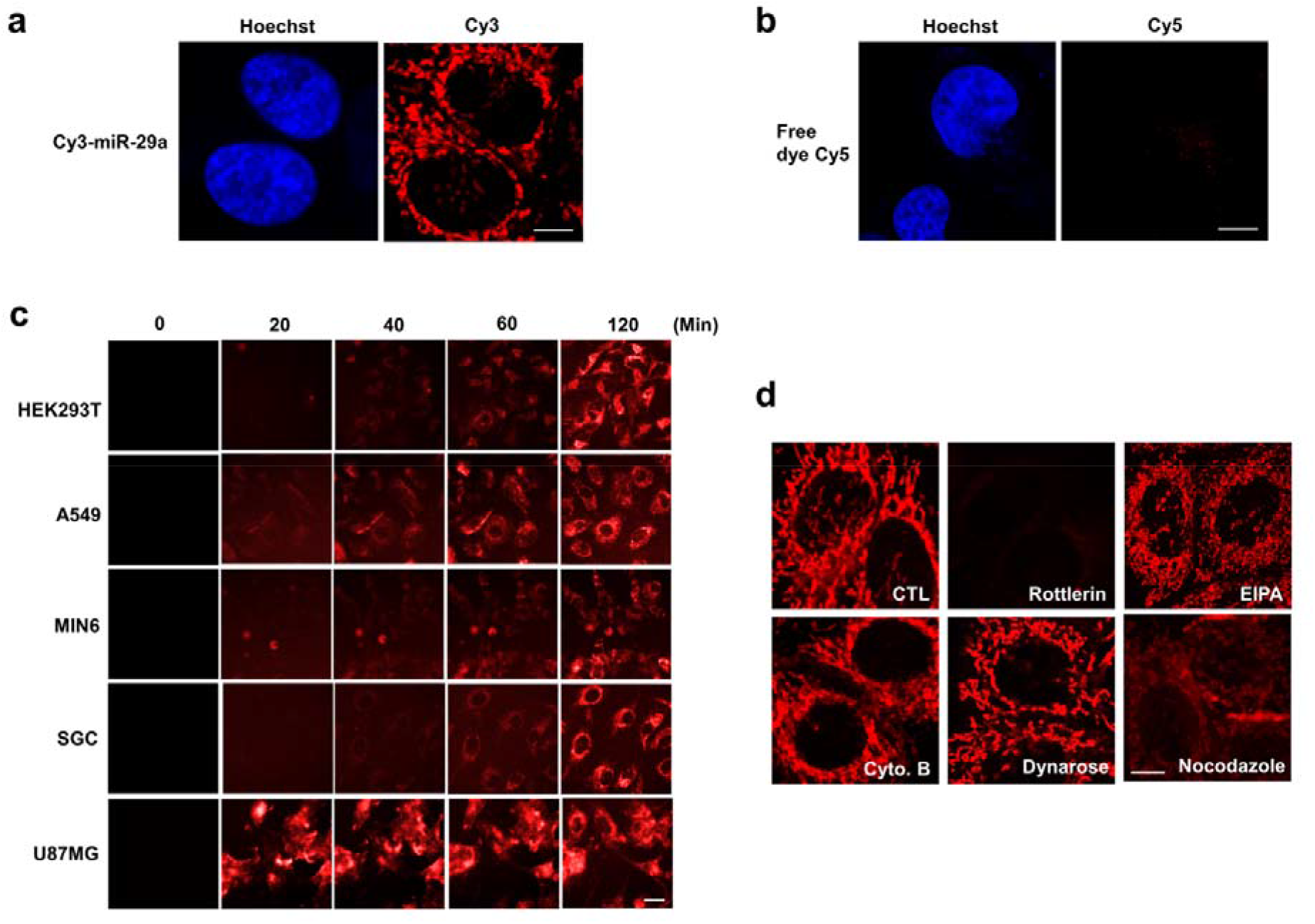
miRNA uptake by various cells. **a**, Uptake of 5’-Cy3-miR-29a by HeLa cells. **b**, No uptake of Cy5 alone by HeLa cells. Scale bars, 10μm. **c**, The time-course of 5’-Cy5-miR-29a uptake by various cells. Scale bars, 20μm. **d**, Effect of cytoskeleton-disrupting reagents on 5’-Cy5-miR-29a uptake by HeLa cells. Scale bars, 10μm.

**Extended Data | Figure 2.**
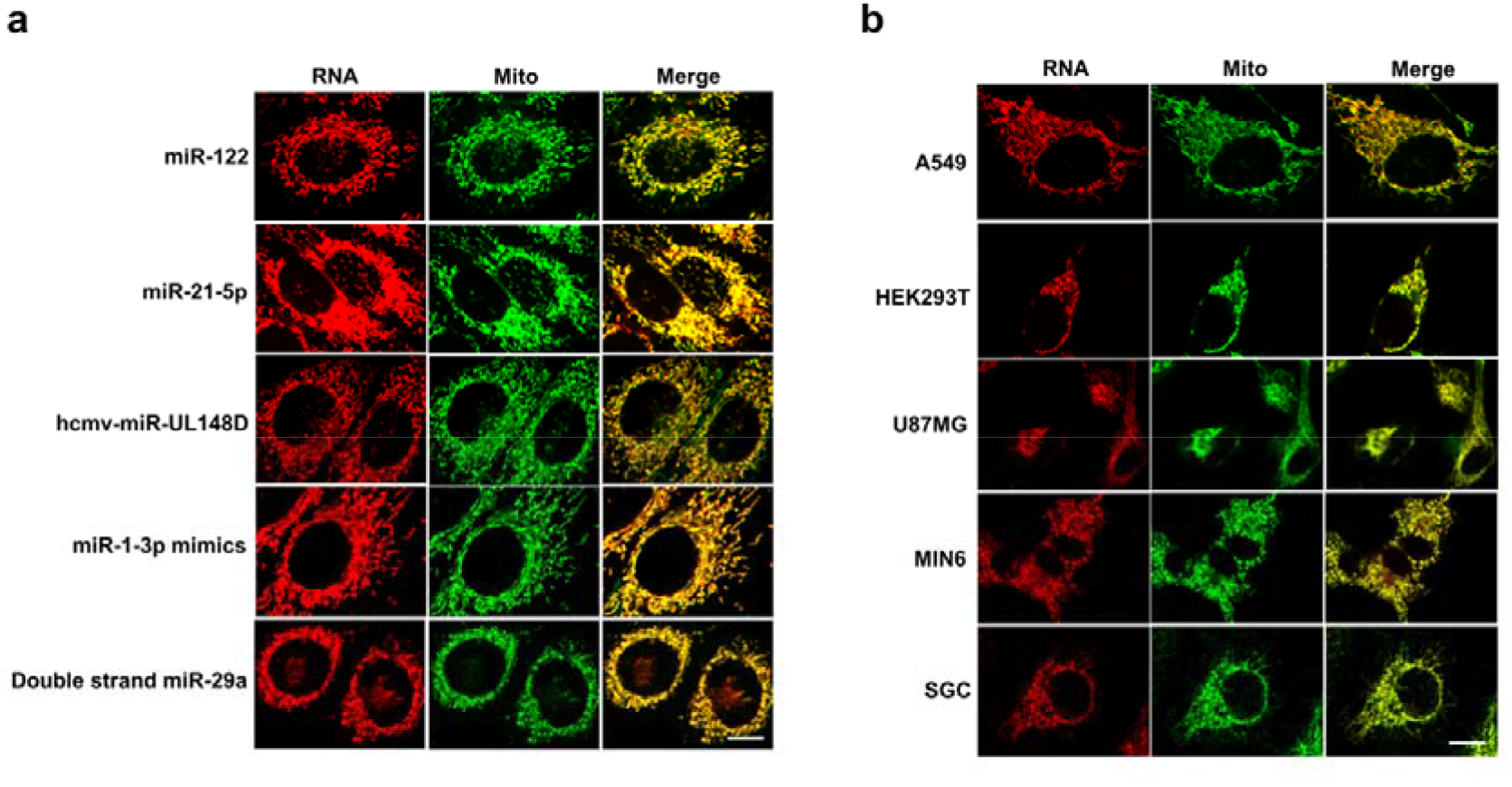
Mitochondria are destination of the internalized miRNAs in various cells. **a**, Colocalization of various structures of internalized small RNAs with mitochondria in HeLa cells after 2 h incubation. **b**, Colocalization of 5’-Cy5-miR-29a with mitochondria in different cells. Scale bars, 10μm.

**Extended Data | Figure 3.**
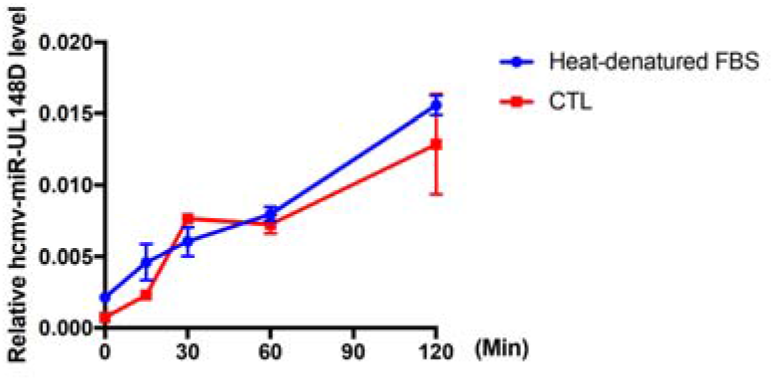
Uptake of hcmv-miR-UL148D by HeLa cells in the presence of 10% control (CTL) or heat-denatured FBS in 120 min.

**Extended Data | Figure 4.**
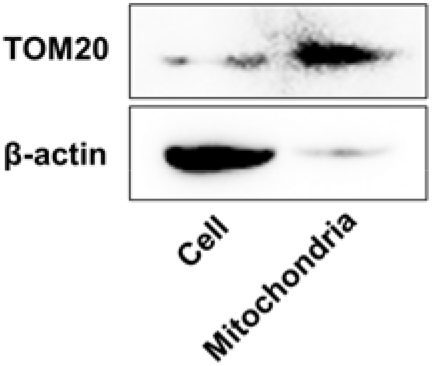
Fraction and identification of mitochondria by western blot analysis.

